# GEM: A manifold learning based framework for reconstructing spatial organizations of chromosomes

**DOI:** 10.1101/161208

**Authors:** Guangxiang Zhu, Wenxuan Deng, Hailin Hu, Rui Ma, Sai Zhang, Jinglin Yang, Jian Peng, Tommy Kaplan, Jianyang Zeng

## Abstract

Decoding the spatial organizations of chromosomes has crucial implications for studying eukaryotic gene regulation. Recently, Chromosomal conformation capture based technologies, such as Hi-C, have been widely used to uncover the interaction frequencies of genomic loci in high-throughput and genome-wide manner and provide new insights into the folding of three-dimensional (3D) genome structure. In this paper, we develop a novel manifold learning framework, called GEM (**G**enomic organization reconstructor based on conformational **E**nergy and **M**anifold learning), to elucidate the underlying 3D spatial organizations of chromosomes from Hi-C data. Unlike previous chromatin structure reconstruction methods, which explicitly assume specific relationships between Hi-C interaction frequencies and spatial distances between distal genomic loci, GEM is able to reconstruct an ensemble of chromatin conformations by directly embedding the neighboring affinities from Hi-C space into 3D Euclidean space based on a manifold learning strategy that considers both the fitness of Hi-C data and the biophysical feasibility of the modeled structures, which are measured by the conformational energy derived from our current biophysical knowledge about the 3D polymer model. Extensive validation tests on both simulated interaction frequency data and experimental Hi-C data of yeast and human demonstrated that GEM not only greatly outperformed other state-of-art modeling methods but also reconstructed accurate chromatin structures that agreed well with the hold-out or independent Hi-C data and sparse geometric restraints derived from the previous fluorescence *in situ* hybridization (FISH) studies. In addition, as GEM can generate accurate spatial organizations of chromosomes by integrating both experimentally-derived spatial contacts and conformational energy, we for the first time extended our modeling method to recover long-range genomic interactions that are missing from the original Hi-C data. All these results indicated that GEM can provide a physically and physiologically valid 3D representations of the organizations of chromosomes and thus serve as an effective and useful genome structure reconstructor.

## Introduction

The three-dimensional (3D) organizations of chromosomes in nucleus are closely related to diverse genomic functions, such as transcription regulation, DNA replication and genome integrity [1–4]. Therefore, decoding the 3D genomic architecture has important implications in revealing the underlying mechanisms of gene activities. Unfortunately, our current understanding on the 3D genome folding and the related cellular functions still remains largely limited. In recent years, the proximity ligation based chromosome conformation capture (3C) [5, 6], and its extended methods, such as Hi-C [7] and chromatin interaction analysis by paired-end tag sequencing (ChIA-PET) [8], have provided a revolutionary tool to study the 3D organizations of chromosomes at different resolutions in various cell types, organisms and species by measuring the interaction frequencies between genomic loci nearby in space.

To gain better mechanistic insights into understanding the 3D folding of the genome, it is necessary to reconstruct the 3D spatial arrangements of chromosomes based on the interaction frequencies derived from 3C-based data. Indeed, the modeling results of 3D genome structure can shed light on the relationship between complex chromatin structure and its regulatory functions in controlling genomic activities[1–4]. However, the modeling of 3D chromatin structure is not a trivial task, as it is often complicated by uncertainty and sparsity in experimental data, as well as high dynamics and stochasticity of chromatin structure itself. Generally speaking, in the 3D genome structure modeling problem, we are given Hi-C data, which can be represented by a matrix where each element represents the interaction frequency of a pair of genomic loci, and our goal is to reconstruct the 3D organization of genome structure and obtain the 3D spatial coordinates of all genomic loci. In practice, in addition to Hi-C data, additional known constraints, such as shape and size of nucleus, can also be integrated to achieve more reliable modeling results and further enhance the physical and biological relevance of the reconstructed genomic structure [9, 10].

In recent years, numerous computational methods have been developed to reconstruct the 3D organizations of chromosomes [5, 7, 11–28]. Most of these approaches, such as the multi-dimensional scaling (MDS) based method [29], ChromSDE [17], ShRec3D [18] and miniMDS [27], heavily depended on the formula *F* ∝ 1/*D*^α^ to represent the conversion from interaction frequencies *F* to spatial distances *D* (where α is a constant). Instead of using the above relationship of inverse proportion, BACH [16] employed a Poisson distribution to define the relation between Hi-C interaction frequencies, spatial distances and other genomic features (e.g., fragment length, GC content and mappability score). After converting Hi-C interaction frequencies into distances, these previous modeling approaches applied various strategies to reconstruct chromatin organizations that satisfy the derived distance constraints. Among them, the optimization based methods, such as the MDS based model [29] and ChromSDE [17], formulated the 3D chromatin structure modeling task into a multivariate optimization problem which aims to maximize the agreement between the reconstructed structures and the distance constraints derived from Hi-C interaction frequencies. More specifically, the MDS based method [29] minimized a strain or stress functions [30] describing the level of violation in the input distance constraints, while ChromSDE [17] used a semi-definite programming technique to elucidate the 3D chromatin structures. In [19], an expectation-maximization based algorithm was proposed to infer the 3D chromatin organizations under a Bayesian inference framework. Several stochastic sampling based methods, such as Markov chain Monte Carlo (MCMC) and simulated annealing [31], were also used in a probabilistic framework to compute chromatin structures that satisfy the spatial distances derived from Hi-C data. In addition, a shortest-path algorithm was used in ShRec3D [18] to interpolate the spatial distance matrix obtained from Hi-C data, based on which the MDS algorithm was then applied to reconstruct the 3D coordinates of genomic loci.

Despite the significant progress made in the methodology development of 3D chromatin structure reconstruction, most of existing reconstruction methods still suffer from several limitations. For example, few methods integrate the experimental Hi-C data with the previously known biophysical energy model of 3D chromatin structure, raising potential concerns about the biophysical feasibility and structural stability of the reconstructed 3D structures. More importantly, as mentioned previously, most of existing chromatin structure modeling methods [5, 7, 11, 13, 15–24, 26, 27] heavily rely on the underlying assumptions about the explicit relationships between interaction frequencies derived from 3C-based data and spatial distances between genomic loci, which may cause bias during the modeling process.

Recently, manifold learning, such as t-SNE [32], has been successfully applied as a general framework for nonlinear dimensionality reduction in machine learning and pattern recognition [30, 33–35]. It aims to reconstruct the underlying low-dimensional manifolds from the abstract representations in the high-dimensional space. In this work, to address the aforementioned issues in 3D chromatin structure reconstruction, we propose a novel manifold learning based framework, called GEM (**G**enomic organization reconstructor based on conformational **E**energy and **M**anifold learning), which directly embeds the neighboring affinities from Hi-C space into 3D Euclidean space using an optimization process that considers both Hi-C data and the conformational energy derived from our current biophysical knowledge about the polymer model. From the perspective of manifold learning, the spatial organizations of chromosomes can be interpreted as the geometry of manifolds in 3D Euclidean space. Here, the Hi-C interaction frequency data can be regarded as a specific representation of the neighboring affinities reflecting the spatial arrangements of genomic loci, which is intrinsically determined by the underlying manifolds embedded in Hi-C space. Based on this rationale, manifold learning can be applied here to uncover the intrinsic 3D geometry of the underlying manifolds from Hi-C data.

Our extensive tests on both simulated and experimental Hi-C data [7, 14] showed that GEM greatly outperformed other state-of-start modeling methods, such as the MDS based model [29], BACH [16], ChromSDE [17] and ShRec3D [18]. In addition, the 3D chromatin structures generated by GEM were also consistent with the distance constrains driven from the previously known fluorescence *in situ* hybridization (FISH) imaging studies [36, 37], which further validated the reliability of our method. More intriguingly, the GEM framework did not make any explicit assumption on the relationship between interaction frequencies derived from Hi-C data and spatial distances between genomic loci, and instead it can accurately and objectively infer the latent function between them by comparing the modeled structures with the original Hi-C data.

Considering the dynamic nature of chromatin structures [2, 38, 39], we model the chromatin structures by an ensemble of conformations (i.e., multiple conformations with mixing proportions) instead of a single conformation. Furthermore, as a novel extended application of the GEM framework, we have introduced a structure-based approach to recover the long-range genomic interactions missing in the original Hi-C data mainly due to experimental uncertainty. We demonstrated this new application of our chromatin structure reconstruction method on both Hi-C and capture Hi-C data, and showed that the recovered distal genomic contacts can be well validated through different interaction frequency datasets or epigenetic features. The competence to recover the missing long-range genomic interactions not only offers a novel application of GEM but also provides a strong evidence indicating that GEM can yield a physically and physiologically reasonable representation of the 3D organizations of chromosomes.

## Results

### Overview of the GEM framework

We introduced a novel modeling method, called GEM (**G**enomic organization reconstructor based on conformational **E**nergy and **M**anifold learning), to reconstruct the three-dimensional (3D) spatial organizations of chromosomes from the 3C-based interaction frequency data. In our modeling framework, each chromatin structure is considered a linear polymer model, i.e., a consecutive line consisting of individual genomic segments. In particular, each restriction site cleaved by the restriction enzyme is abstracted as an end point (which we will also refer to as a node or genomic locus) of a genomic segment and the line connecting every two consecutive end points represents the corresponding chromatin segment between two restriction sites. This model has been widely used as an efficient and reasonably accurate model given the current resolution of Hi-C data [15–19].

In the GEM pipeline (Fig. 1), we first model the input Hi-C interaction frequency data as a representation of neighboring affinities between genomic loci in Hi-C space, and then construct an interaction network (in which each edge indicates an interaction frequency between two genomic loci) to reflect the organizations of chromosomes in Hi-C space. Our goal is to embed the organizations of chromosomes from Hi-C space into 3D Euclidean space such that the embedded structures preserve the neighborhood information of genomic loci, while also maintaining the stable structures as possible (i.e., with the minimum conformational energy). The meaningful spatial organizations of chromosomes can be interpreted as the geometry of manifolds in 3D Euclidean space, while the Hi-C interaction frequency data, i.e., a specific representation of the neighboring affinities reflecting the spatial arrangements of genomic loci, which is intrinsically determined by the underlying manifolds embedded in Hi-C space. Inspired by manifold learning (see “Methods” Supplementary Fig. 1), GEM reconstructs the chromatin structures by directly embedding the neighboring affinities from Hi-C space into 3D Euclidean space using an optimization process that considers both the fitness of Hi-C data and the biophysical feasibility of the modeled structures measured in terms of conformational energy (which is derived mainly based on our current biophysical knowledge about the 3D polymer model). Unlike most of existing methods for modeling chromatin structures from Hi-C data, GEM does not assume any specific relationship between Hi-C interaction frequencies and spatial distances between genomic loci. On the other hand, such a latent relationship can be inferred based on the input Hi-C data and the final structures modeled by GEM.

**Figure 1:**
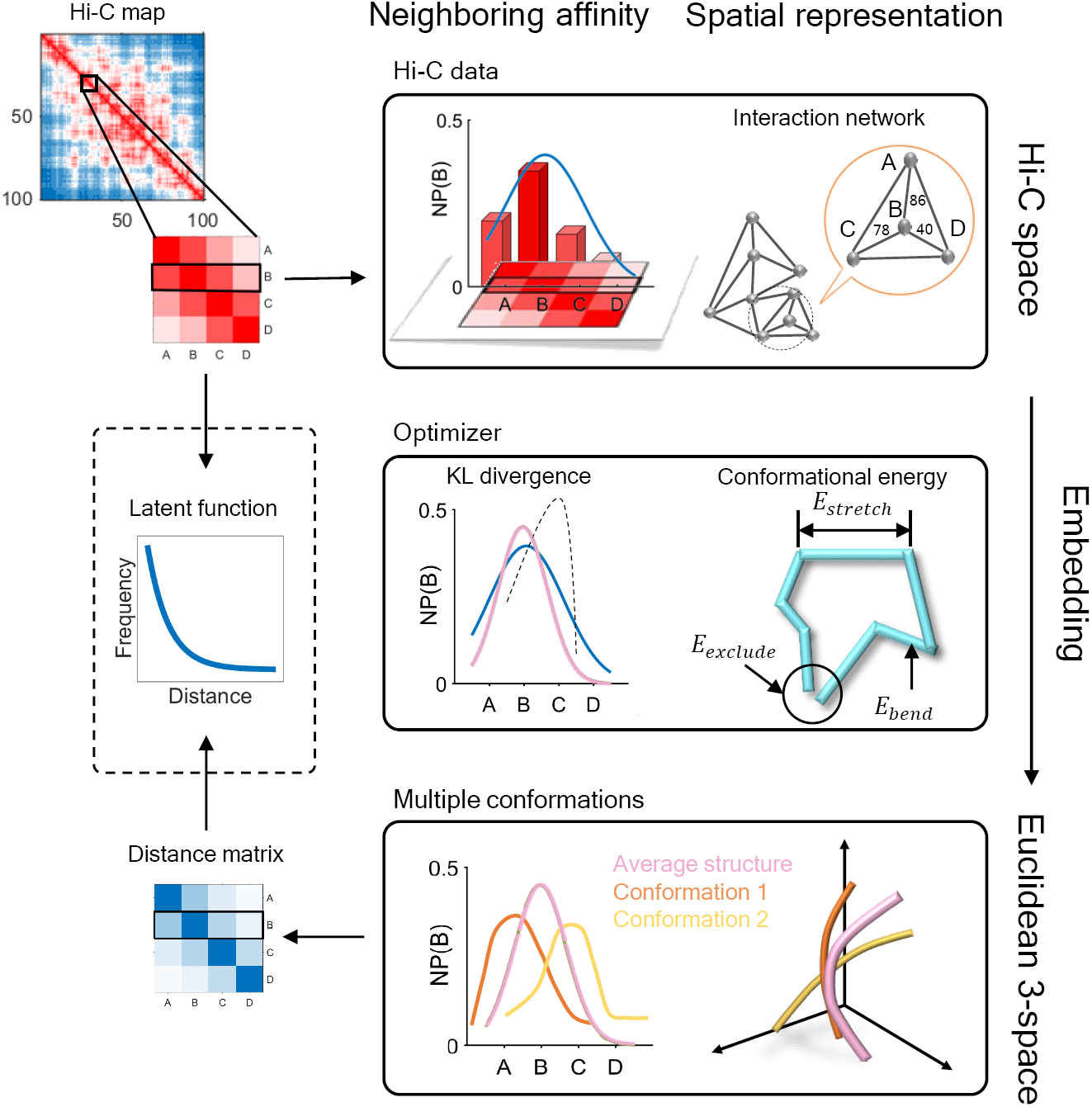
A schematic illustration of the GEM pipeline. The genomic loci A, B, C and D are selected as an example to demonstrate our pipeline. We first build up an interaction network from the input Hi-C data to represent the organizations of chromatin structures in Hi-C space. In this interaction network, each node represents a genomic loci and each edge represents a pairwise interaction describing the neighbouring affinity between genomic loci in Hi-C space. Based on an optimization that considers both the KL divergence between experimental and reconstructed Hi-C data and the conformational energy, the interaction network is then embedded into 3D Euclidean space to reconstruct the 3D chromatin structures. During the embedding process, we first calculate an average conformation as an initial structure, and then refine the initial structure to obtain an ensemble of conformations through a multi-conformation optimization technique (see “Methods”). Finally, we can infer the latent function between Hi-C interaction frequencies and spatial distances between genomic loci based on the input interaction frequency matrix and the output spatial distance matrix derived from GEM (shown in the dashed box). Neighbouring probability, NP(B), in the figure represents the probability of the spatial interaction between current genomic and genomic locus B.

We use *ψ_i_* to represent the *i*-th genomic locus of the chromatin structure Ψ in Hi-C space. Given two genomic loci *ψ*_*i*_ and *ψ*_*j*_, their neighboring affinity, denoted by *p*_*I,j*_, is defined as

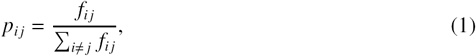

where *f*_*ij*_ stands for the interaction frequency between *ψ*_*i*_ and *ψ*_*j*_. Here, the neighboring affinity represents the probability that two genomic loci are neighbor. The neighborhood of a genomic locus thus can be featured by its neighboring affinities of this genomic locus. Here, we use the normalized interaction frequencies instead of the raw count information, which is more robust and happens to be the same as in t-SNE [32]. Inspired by the idea of t-SNE, we map the Hi-C space representation of a chromosome, denoted by Ψ = {*ψ*_1_, *ψ*_2_, …, *ψ*_*n*_} (where *n* is the total number of genomic loci) into 3D Euclidean space to derive the final 3D chromatin structure, denoted by *S* = {*s*_1_, *s*_2_, …, *s*_*n*_}, where *s*_*i*_ represents the coordinates of the *i*-th genomic locus in 3D Euclidean space, based on a neighboring affinity embedding process, which preserves the neighborhood information of genomic loci in Hi-C space as much as possible. That is, if two genomic loci are neighbor in Hi-C space, they would have a large probability of being neighbor in 3D Euclidean space.

In the t-SNE framework, which is a typical model of manifold learning, a Student t-distribution which generally has much heavier tails than Gaussian distribution is used to alleviate the “crowding problem” (i.e., many close-by neighbors would be placed far off because of limited room when arranging high-dimensional data into low-dimensional space) in the embedding from high-dimensional to low-dimensional space [32]. In our chromatin structure modeling problem, we use *q*_*ij*_ to denote the probability that genomic loci *s*_*i*_ and *s* _*j*_ pick each other as neighbors in 3D Euclidean space after embedding, which is defined a

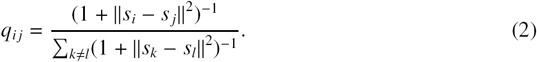

where ∥· ∥ stands for the Euclidean distance.

Chromatin can change dynamically in the nucleus especially during interphase. Thus unlikely its structure can be accurately described by one single consensus conformation. In our framework, we develop a multi-conformation version of the embedding approach to model an ensemble of chromatin conformations. In particular, we use multiple 3D conformations with mixing proportions instead of a single conformation to interpret the Hi-C data. Here, we redefine the joint probability *q*_*ij*_ a

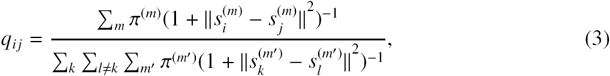

where π^(*m*)^ stands for the mixing proportion of the *m*-th conformation, and 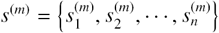 represents the coordinates of the *m*-th conformation.

From the perspective of neighbor embedding [32, 40], if an ensemble of chromatin conformations in 3D Euclidean space, denoted by {(*s*^(1)^, π^(1)^), (*s*^(2)^, π^(2)^), …, (*s*^(*m*)^, π^(*m*)^)}, correctly models the neighborhood system of Ψ in Hi-C space, the joint probabilities *p*_*ij*_ and *q*_*ij*_ should match to each other. As in other t-SNE based learning tasks [41, 42], we minimize the Kullback-Leibler divergence (KL divergence) to find a low-dimensional (3D Euclidean space in our case) data representation that has the lowest degree of mismatch to the original Hi-C data (which can be considered in high-dimensional space). We use *P*_*i*_ to denote the neighborhood system of ψ_*i*_ in Hi-C space and Q _*i*_ to denote the neighborhood system of *s*_*i*_ in 3D Euclidean space. Moreover, we add a conformational energy term *C*_2_ (see “Methods”) to ensure that the modeled structures have high energy stability. That is, the overall cost function *C* is defined as

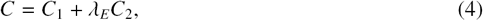

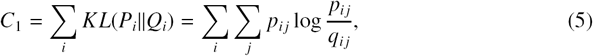

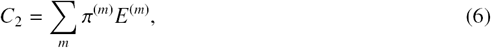

where *E*^(*m*)^ stands for the conformational energy of the *m*-th conformation in the ensemble and *λ_E_* stands for the coefficient that weighs the relative importance between the data term representing the fitness of Hi-C data and the energy term. More details about the optimization of the above cost function *C* can be found in “Methods”.

Taking a deeper look at *C*_1_, it is obvious that KL divergence is not symmetric [42]. From a different perspective, 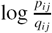 represents a mismatch term and *p*_*ij*_ can be regarded as the weighting factor of such a mismatch term. This observation means that, there is of relatively large cost to use points far from each other (i.e., with small *q*_*ij*_) in 3D Euclidean space to represent nearby genomic loci (i.e., with large *p*_*ij*_) in Hi-C space, while it is of relatively small cost to use nearby points to represent two genomic loci far away in Hi-C space. In other words, GEM aims to preserve local structure when mapping from Hi-C space into 3D Euclidean space. This merit of retaining local structure particularly meets the requirement of 3D chromatin structure modeling, as Hi-C data exactly reflect the topological properties of local structures of chromosomes. In addition, it is reasonable to associate pairs of genomic loci with higher interaction frequencies with more confidence during the modeling process.

### Validation on simulated Hi-C data

We first validated the modeling performance of GEM on the simulated Hi-C data (see “Methods”). The simulated Hi-C data were then fed into GEM to reconstruct the chromatin structures. We tested GEM on different simulated Hi-C maps which were generated by varying a wide ranges of parameter settings during the simulation process (Fig. 2, Supplementary Figs 2-4). Here, we evaluated the Pearson correlations between the distance matrices of our reconstructed models and the original conformations that were used to generate the simulated data. We also compared the modeling performance of GEM to that of three other reconstruction methods, including the MDS based model [29], ChromSDE [17] and ShRec3D [18]. As our simulation process did not consider the sequence content (e.g., GC content) of chromatin structures, here we did not include BACH [16] in the comparison tests on simulated Hi-C data. All the validation tests on synthetic Hi-C data generated by a variety of conditions showed that GEM achieve the best modeling performance in terms of the closeness to the original structures that were used to generate the simulated data (Fig. 2a, Supplementary Figs 2a, 3a and 4a).

**Figure 2:**
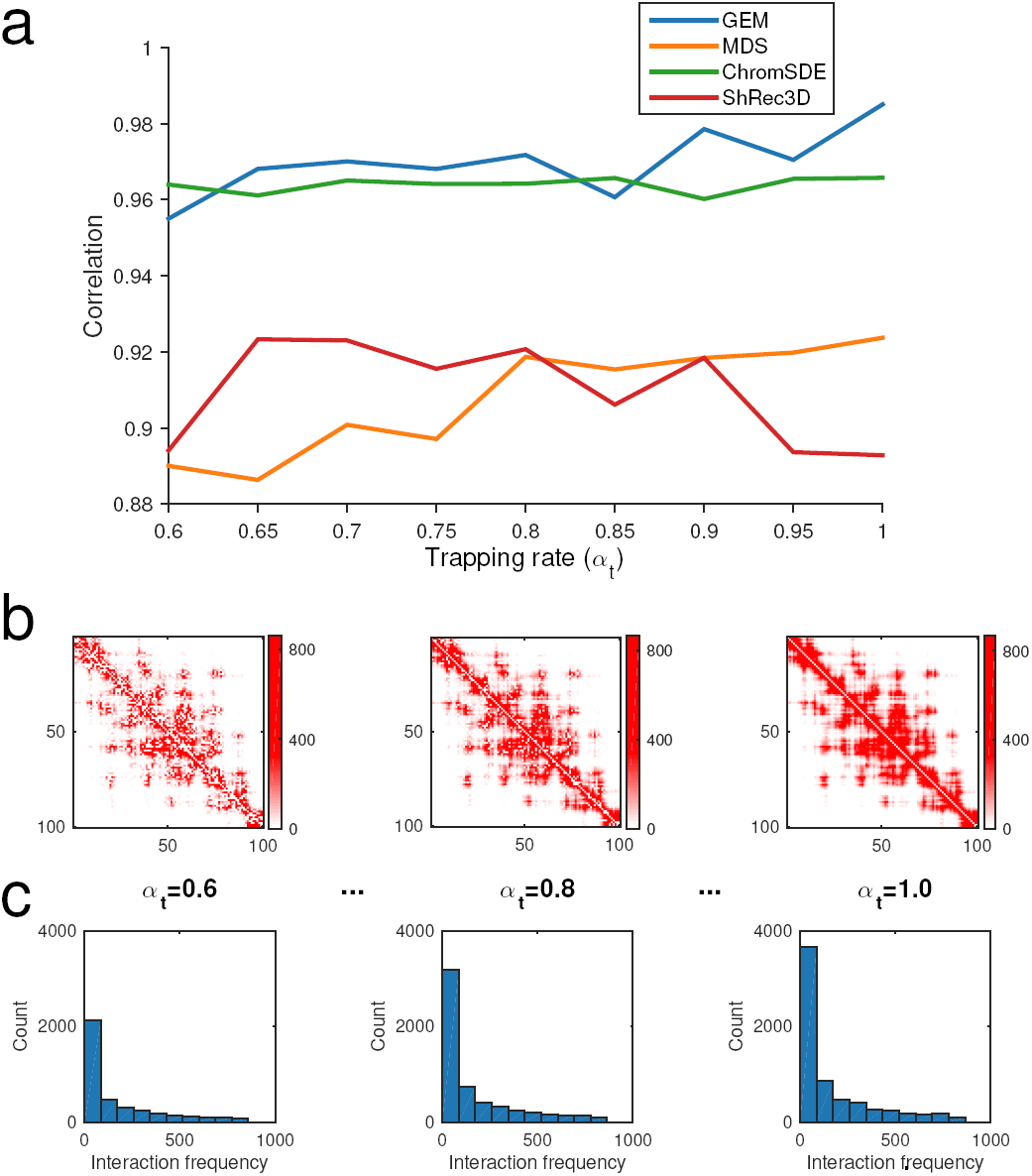
The validation results on the simulated Hi-C data, which were generated according to different settings of the trapping rate *α_t_* (see “Methods”). (**a**) The comparisons of Pearson correlations between GEM and other modeling methods, including the MDS based model [29], ChromSDE [17] and ShRec3D [18]. (**b**) and (**c**) show the typical examples of the simulated Hi-C maps and the corresponding distributions of the reconstructed interaction frequencies as *α_t_* increases, respectively.

### Validation on experimental Hi-C data

We then evaluated the modeling performance of GEM on experimental Hi-C data [7, 14]. We first used Pymol [43] to visualize the overall ensemble of the chromatin conformations reconstructed by GEM, taking human chromosome 14 at a resolution of 1 Mb as an example (Fig. 3a). The modeled 3D organizations of chromosomes can provide a direct and vivid visualization about the 3D spatial arrangements of chromosomes, which may offer useful mechanistic insights about the 3D folding of chromatin structure and its functional roles in gene regulation. Through simple visual inspection of the ensemble of four chromatin conformations reconstructed by GEM (Fig. 3a), we observed that they displayed similar but not identical 3D spatial organizations. In addition, we found that they are all organized into alike obvious isolated regions that agreed well with those identified from the Hi-C map. Such consistency of domain partition also suggested the reasonableness of the chromatin conformations reconstructed by GEM. Also, the similar domain partitions of different conformations were consistent with the previous studies [28, 44] that topological domains are hallmarks of chromosomal conformations in despite of their dynamic structrual variability.

**Figure 3:**
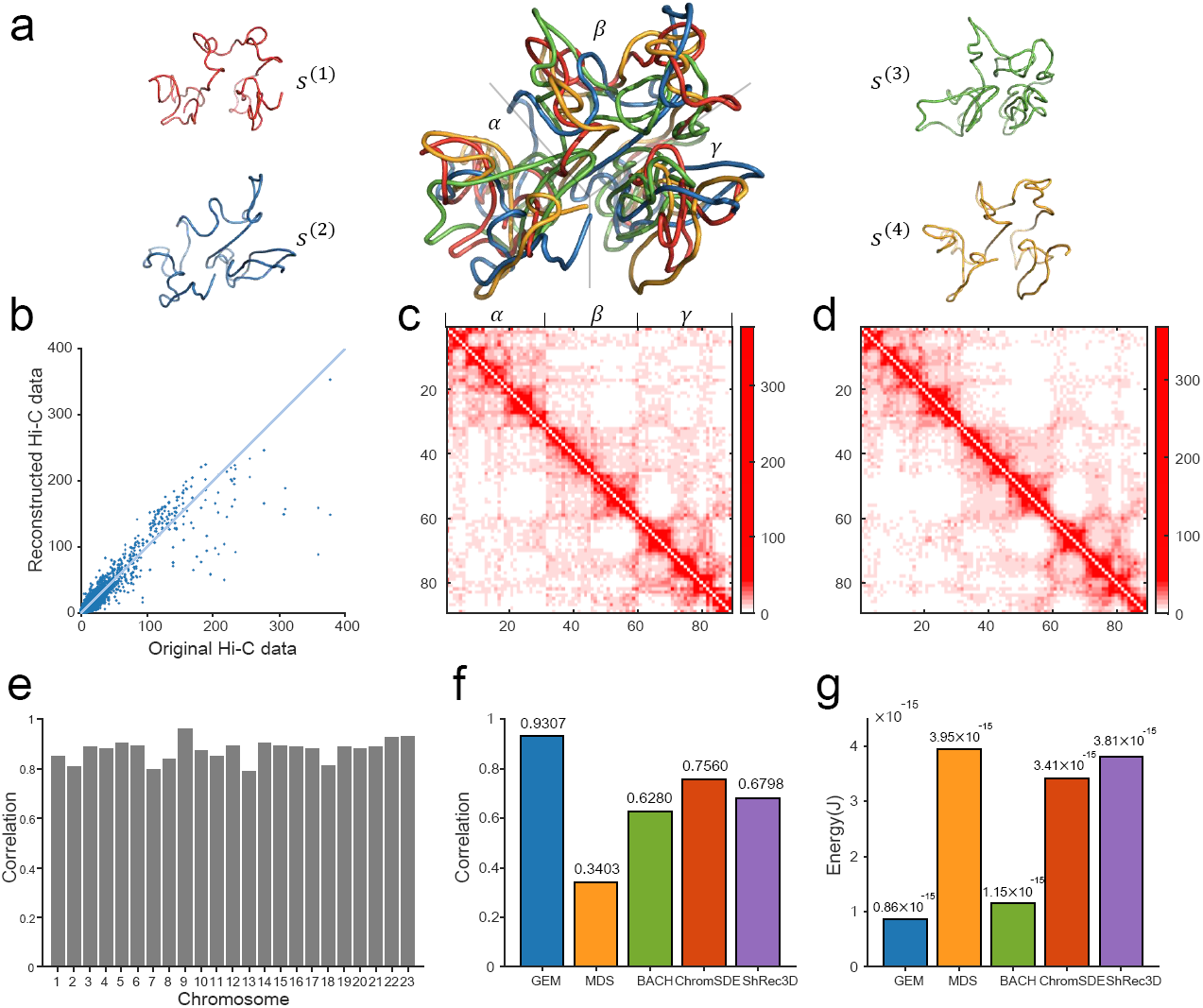
The chromatin structure modeling results on human chromosomes under 1 Mb resolution. (**a**) Visualization of the computed ensemble of human chromosome 14. The four conformations *s*^(1)^, *s*^(2)^, *s*^(3),^ *s*^(4)^ in the derived ensemble are shown in red, blue, green and orange, respectively. The middle shows the superimposition of all four conformations, which were all aligned using the singular value decomposition (SVD) algorithm [61]. The three large isolated regions (*α, β, γ*) which can be facilely distinguished from the reconstructed 3D conformations were consistent well with those detected based on the original Hi-C map (see Panel c). (**b**) The 10-fold cross-validation results for human chromosome 14, in which the scatter plot of the reconstructed Hi-C data derived from the modeled structures vs. the original Hi-C data is shown. (**c**,**d**) The original interaction frequency map derived from experimental Hi-C data and the reconstructed Hi-C map predicted by the modeled structures for human chromosome 14 in the 10-fold cross-validation results, respectively. (**e**) Bar graph depicting mutual validation by two sets of experimental Hi-C data for individual 23 human chromosomes, which were collected using two different restriction enzymes (i.e., HindIII vs. NcoI), respectively. (**f**,**g**) Comparison results between different modeling methods, in terms of the agreement between experimental and predicted Hi-C data and the conformational energy, respectively.

Next, we performed a 10-fold cross-validation procedure to assess the modeling performance of GEM on experimental Hi-C data (see “Methods”). Our 10-fold cross-validation on human Hi-C data demonstrated that GEM was able to reconstruct accurate 3D chromatin structures that agreed well with the hold-out test data. For example, the predicted Hi-C data of human chromosome 14 inferred from the reconstructed conformations were consistent with the original experimental Hi-C data, with the Pearson correlation above 0.93 (Fig. 3b-d).

We also conducted a mutual validation based on different Hi-C datasets collected from distinct experimental platforms. In particular, we chose two Hi-C datasets [7] that were collected using two different restriction enzymes (i.e., HindIII vs. NcoI). These two datasets were fed into GEM separately and their modeling results were then evaluated by cross examining the correlations between the chromatin structures reconstructed from individual datasets. Such a mutual validation indicated that GEM was able to elucidate accurate chromatin structures that were consistent with the other independent dataset, achieving the Pearson correlations of distance matrices close or above 0.8 (Fig. 3e).

In the above cross-validation tests, we also compared the modeling results of GEM to those of other existing methods, including the the MDS based model [29], BACH [16], ChromSDE [17] and ShRec3D [18]. The comparisons demonstrated that GEM outperformed other four modeling methods, in terms of the Pearson correlation between the reconstructed interaction frequency data and the original experimental Hi-C data (Fig. 3f). Here, the reconstructed Hi-C maps for the other modeling methods were computed according to their hypothesis functions (MDS based model, ChromSDE and ShRec3D) or distributions (BACH) on the relationships between interaction frequencies and spatial distances between genomic loci. In addition, since our method also considered the conformational energy term during the modeling process, its reconstructed structures had significantly lower energy than those modeled by other four approaches (Fig. 3g), which implied that GEM can yield biophysically more reasonable spatial representations of the observed Hi-C data.

Taking together, the above validation tests on experimental Hi-C data demonstrated that GEM can outperform other existing modeling methods, and reconstruct an ensemble of more accurate and biophysically more reasonable 3D organizations of chromosomes.

### Validation on FISH data

In addition to the cross-validation tests on experimental Hi-C data, we also verified the modeled chromatin structures using a sparse set of known pairwise distance constraints between genomic loci driven by the FISH imaging technique (Fig. 4). In particular, we examined the agreement between the chromatin structures reconstructed by GEM and the sparse FISH distance constraints obtained from the previously known studies [7, 36, 37], which included ARS603-ARS606, ARS606-ARS607, ARS607-ARS609 on yeast chromosome 6 and L1-L3, L2-L3, L2-L4 on human chromosome 14 (Fig. 4). We compared the average distances between genomic loci driven from the FISH imaging data and our reconstructed models. In addition, we also analyzed the reasonableness of the pairwise spatial distances between genomic loci predicted by GEM based on the relative sequence distances and corresponding compartmentalization information [7]. On yeast chromosome 6 (Fig. 4a), the reconstructed distance between ARS603 and ARS606 was relatively larger than those of other pairs, which was consistent with the fact that the pair ARS603-ARS606 crosses two different compartments (A and B), while the other two pairs (i.e., ARS606-ARS607 and ARS607-ARS609) are located within the same compartment (B). In the same compartment (B), the reconstructed distance between ARS606 and ARS607 was less than that between genomic loci ARS607 and ARS609, suggesting that a pair of genomic loci with small sequence distance preferentially stay close in space, which was also consistent with the previous studies [5, 36]. On human chromosome 14 (Fig. 4b), the reconstructed distance between genomic loci L1 and L3 was notably smaller than that between L2 and L3, which agreed with the fact that, L2 and L3 are closer along the sequence but belong to different compartments (L2 in B and L3 in A), while L1 and L3 are further far away along the sequence but belong to the same compartment (A). These validation results showed that the chromatin structures modeled by GEM were in good agreement with the known pairwise distance constraints derived from FISH data in terms of both average spatial distances and compartment partition, which further verified the modeling power of our method.

**Figure 4:**
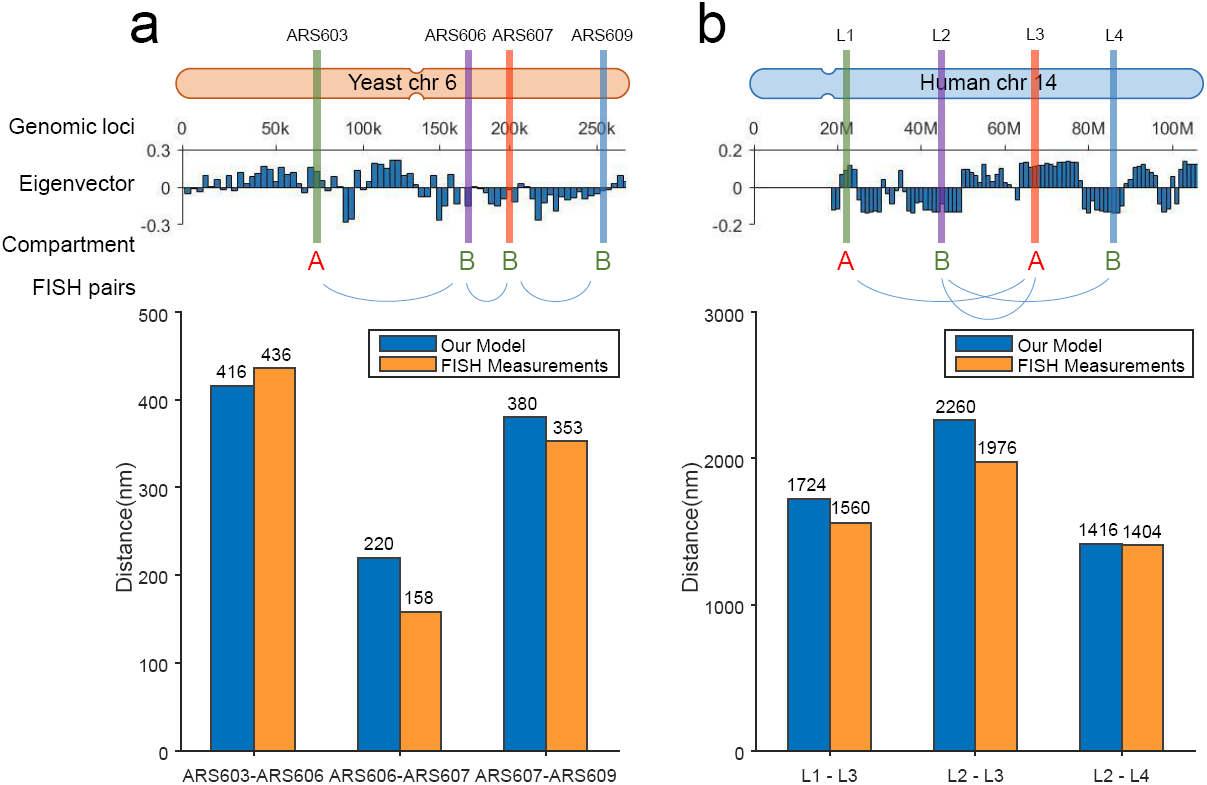
The validation results on the known pairwise distance constraints derived from the FISH imaging data of yeast and human. Top shows the schematic illustrations of the locations of genomic loci used in the validation. Compartment partition was performed based on the eigenvectors of the Hi-C maps computed by principal component analysis (PCA) [7]. Bottom shows the bar graphs depicting the comparisons between the mean distances between genomic loci derived from FISH imaging data and reconstructed by GEM. (**a**) The validation results on the FISH imaging data of yeast chromosome 6. ARS603, ARS606, ARS607 and ARS609 lie consecutively along the chromosome. ARS603 belongs to compartment A, while the other three loci belong to compartment B, as visualized by the schematic illustration (top). (**b**) The validation results on the FISH imaging data of human chromosome 14. L1, L2, L3 and L4 lie consecutively along the chromosome. L1 and L3 belong to compartment A, while L2 and L4 belong to compartment B, as visualized by the schematic illustration (top).

### Analysis of the relationships between Hi-C interaction frequencies and spatial distances

Most of existing chromatin structure modeling approaches assumed that there exists a specific relationship between Hi-C interaction frequencies *F* and the corresponding spatial distances *D* between genomic loci [5, 7, 11, 13, 15–24, 26, 27]. Many prevalent frameworks, such as the MDS based method [29], ChromSDE [17], ShRec3D [18] and miniMDS [27], used the inverse proportion formula *F* ∝ 1/*D*^α.^ The probability based method such as BACH [16] employed Poisson distribution to represent the relationship between Hi-C interaction frequencies, distances and other genomic features (e.g., fragment length, GC content and mappability score). The modeling process of these approaches can be basically divided into two stages: First, the Hi-C interaction frequencies were converted into spatial distances based on the specific assumption on their relationships. Second, the 3D chromatin structures were modeled according to the converted spatial distance constraints. If the specific forms of hypothetical functions or distributions between interaction frequencies and spatial distances are not sufficiently accurate, they will mislead the optimization process and cause bias during the modeling process. Thus, the accuracy of the chromatin structures reconstructed by these methods heavily relied on the assumed relationships between interaction frequencies and spatial distances.

Here, we argued that the specific assumptions about the relationships between Hi-C interaction frequencies and spatial distances in most of previous chromatin structure modeling approaches are not advisable, based on our tests on the simulated Hi-C data, in which the true relationships between interaction frequencies and spatial distances were considered known and thus can be used to examine all possible hypothetical functions defining their relationships. First, the latent relationships between interaction frequencies and spatial distances can be affected by various factors and tend to display different concrete forms despite their similar inverse proportion forms. In addition, the relationships are generally complex and it is usually difficult to describe them by a consensus expression. As validated by the simulated Hi-C data (Fig. 5), the latent functions between Hi-C interaction frequencies and spatial distances varied on the simulated Hi-C data generated according to different conditions. Moreover, many previous modeling approaches [14–18] mainly focused on the reciprocal forms (e.g., *F* ∝ 1/*D*^α^) between interaction frequencies and spatial distances and ignored the proportional factor. Thus, the final modeled structures were merely the scaled models of the true conformation conformations. To obtain the exact true structures, knowledge about the scaling factors was also required in these modeling methods.

**Figure 5:**
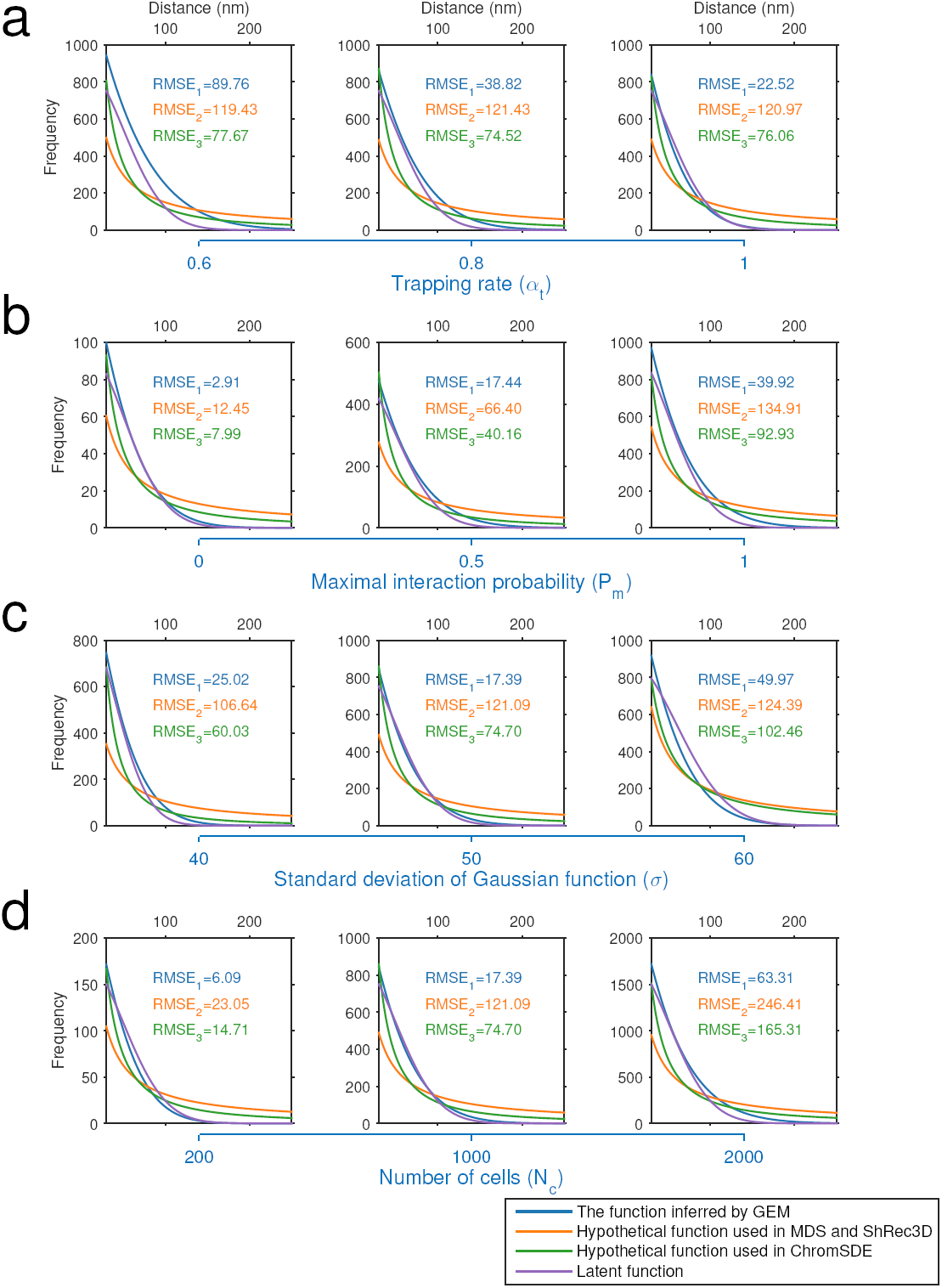
Relationships between Hi-C interaction frequencies and reconstructed spatial distances derived based on different test settings of simulated Hi-C data. The purple curves depict the latent relationships between Hi-C interaction frequencies and reconstructed spatial distances derived based on the tests on simulated Hi-C data, which were generated according to different settings of the trapping rate *α_t_* (**a**), the maximum interaction probability *P*_*m*_ (**b**), the standard deviation of Gaussian function σ (**c**), and the number of cells *N*_*c*_ (**d**), respectively. The blue, orange and green curves show the functions inferred by GEM, the hypothetical function *F α* 1/*D* used in the MDS based model [29] and ShRec3D [18], and the hypothetical function *F α* 1/*D*^α^ used in ChromSDE [17], respectively. The root-mean-square error (RMSE) was used to measure the distances between these functions used in the modeling frameworks (shown in blue, orange or green curves) and the latent functions (shown in purple curves), which can be derived from the parameter settings used to generate the simulated Hi-C data.

To overcome the aforementioned drawbacks in the previous chromatin structure modeling methods, we developed a novel manifold learning based framework to reconstruct the 3D spatial organizations of chromatin that does not require any pre-assumed relationship or function between Hi-C interaction frequencies and spatial distances (Fig. 1). In particular, we directly obtained the 3D chromatin structures by embedding the neighboring affinities from Hi-C space into 3D Euclidean space through a manifold learning based strategy (see “Methods”). In addition, by taking both the fitness of Hi-C data and the structure stability measured in terms of conformational energy into consideration during the embedding process, the reconstructed chromatin structures can also hold the true scale.

By comparing the modeled structures with the original Hi-C data (as shown in the dashed box in Fig. 1), GEM can also infer the latent relationships between interaction frequencies and spatial distances. We used the tests on simulated Hi-C data to demonstrate this point. Specifically, the plotted scatters between the simulated Hi-C interaction frequencies and the spatial distances derived from our modeled structures showed that there existed a certain function between them, which was roughly in a reciprocal form that had been widely accepted in the literature of chromatin structure modeling [7, 15, 45]. We further estimated the latent functions in more detail by curve fitting into the scatter plots (which is implemented by finding the proper function forms and parameters with the lowest RMSEs to interpret the scatters; Fig. 5). The comparisons showed that our derived expressions were much closer to the real functions (which can be obtained from the simulated Hi-C data) between Hi-C interaction frequencies and spatial distances than the specific inverse proportion formulas assumed in the previous modeling approaches, including the MDS based model [29], ShRec3D [18] and ChromSDE [17] (Fig. 5). These results suggested that GEM can accurately capture the latent relationships between Hi-C interaction frequencies and spatial distances without making any specific assumption on the specific forms of their inverse proportion relationships during the structure modeling process. Next, we analyzed the derived relationships between Hi-C interaction frequencies and spatial distances reconstructed by GEM on experimental Hi-C data (Fig. 6). Indeed, there existed a certain inverse proportion function between the experimental Hi-C interaction frequencies and the reconstructed spatial distances, which can be confirmed by the goodness of the fitting results measured by the root-mean-square errors (RMSEs). In addition, our investigation showed that chromatin structures from different chromosomes, at different resolutions or from different species can display distinct inverse proportion forms defining the relationships between Hi-C interaction frequencies and reconstructed spatial distances. This result further implied that it would be generally unadvisable to assume the existence of a single consensus expression for the relationships between Hi-C interaction frequencies and spatial distances between genomic loci.

**Figure 6:**
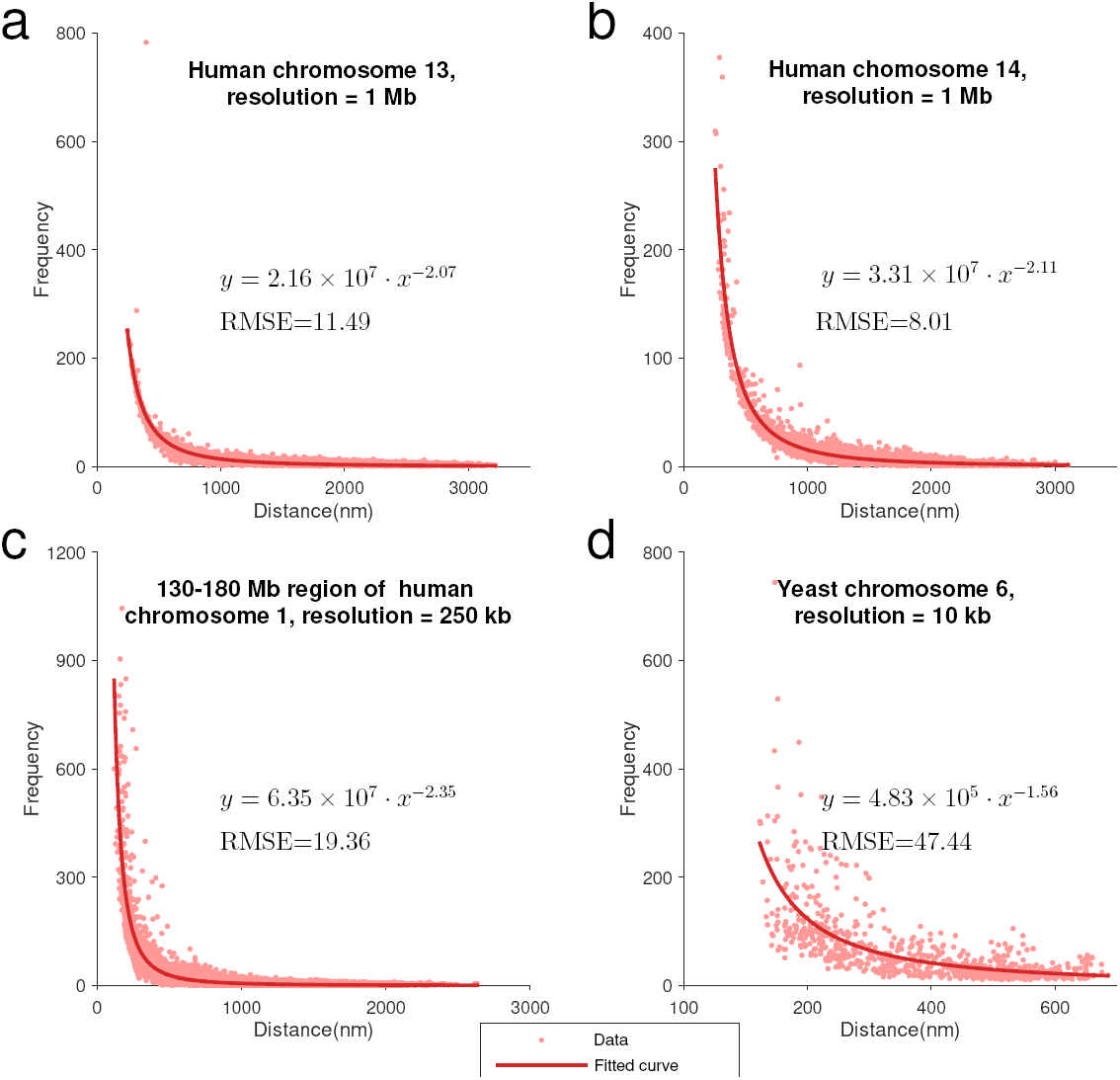
Relationships between Hi-C interaction frequencies and reconstructed spatial distances derived from the chromatin structures modeled by GEM on experimental Hi-C data. (**a**-**d**) The latent functions inferred by GEM between Hi-C interaction frequencies and reconstructed spatial distances on human chromosome 13 at 1Mb resolution, human chromosome 14 at 1Mb resolution, a 130Mb-180Mb region of human chromosome 1 at 250 kb resolution, and yeast chromosome 6 at 10 kb resolution, respectively. The functions were obtained by curve fitting to the points representing the pairs of Hi-C interaction frequencies and reconstructed spatial distances in the modeled structures. The expressions of the derived functions and the fitting results measured in terms of the root-mean-square errors (RMSEs) are also shown.

### Application of the modeled chromatin structures to recover missing long-range genomic interactions

Most of previous studies mainly used the 3D chromatin structures reconstructed from Hi-C data to visualize and inspect the topological and spatial arrangements among different genomic regions [5, 7, 11–28]. The modeled chromatin structures were rarely applied to expand the geometric constraints derived from the original experimental Hi-C data. On the other hand, due to experimental uncertainty, Hi-C data may miss a certain number of long-range genomic interactions or contain extra noisy spatial contacts between distal genomic loci. Nevertheless, the long-range spatial contacts derived from current Hi-C data are generally able to provide a sufficient number of geometric restraints to reconstruct accurate 3D scaffolds of chromosomes. In addition, the conformational energy incorporated in our modeling framework can provide an extra type of restraints to infer biophysically stable and reasonable chromatin structures. For example, conformational energy can provide useful information about the stretching and bending conditions of the chromatin fibres. Thus, the 3D chromatin scaffolds derived by GEM can provide accurate chromatin structure templates to recover those long-range genomic interactions that were missing in the original Hi-C map. This potential application can also be supported by the previous excellent validation results of GEM (Fig. 3). For example, the 10-fold cross-validation results showed that the reconstructed Hi-C map inferred from the reconstructed conformations derived by GEM was consistent with the hold-out dataset in the original experimental data (Fig. 3b-d), which basically indicated that the reconstructed structures can also be used to restore the missing long-range genomic interactions from the original input Hi-C data. Also, the additional validation tests on cross-platform Hi-C data demonstrated that the 3D chromatin conformations reconstructed by GEM from one Hi-C dataset can fit well into another independent dataset (Fig. 3e).

We further used the tests on the Hi-C data [46] collected from different replicates or platforms to demonstrate the potential application of GEM in the recovery of the missing long-range genomic interactions in the original Hi-C map. We first fed the Hi-C data of one replicate into GEM and then used the Hi-C map from the other replicate to validate the missing loops indicated by the modeled chromatin structures. In particular, we looked into the fraction of missing distal chromatin loops that can be validated through another independent dataset. We found that the missing chromatin loops detected by GEM exhibited much closer spatial contacts than the background (i.e., all the reconstructed distances; rank sum test, *P* < 1 × 10^-23^; Fig. 7a,b). In addition, the distributions of the reconstructed distances of missing and known loops (which were present in the Hi-C data of current replicate) were actually close to each other, with the probability plot correlation coefficients above 0.97 (Fig. 7a,b). These observations implies that the missing chromatin loops in Hi-C maps can be potentially restored by the chromatin structures modeled by GEM.

**Figure 7:**
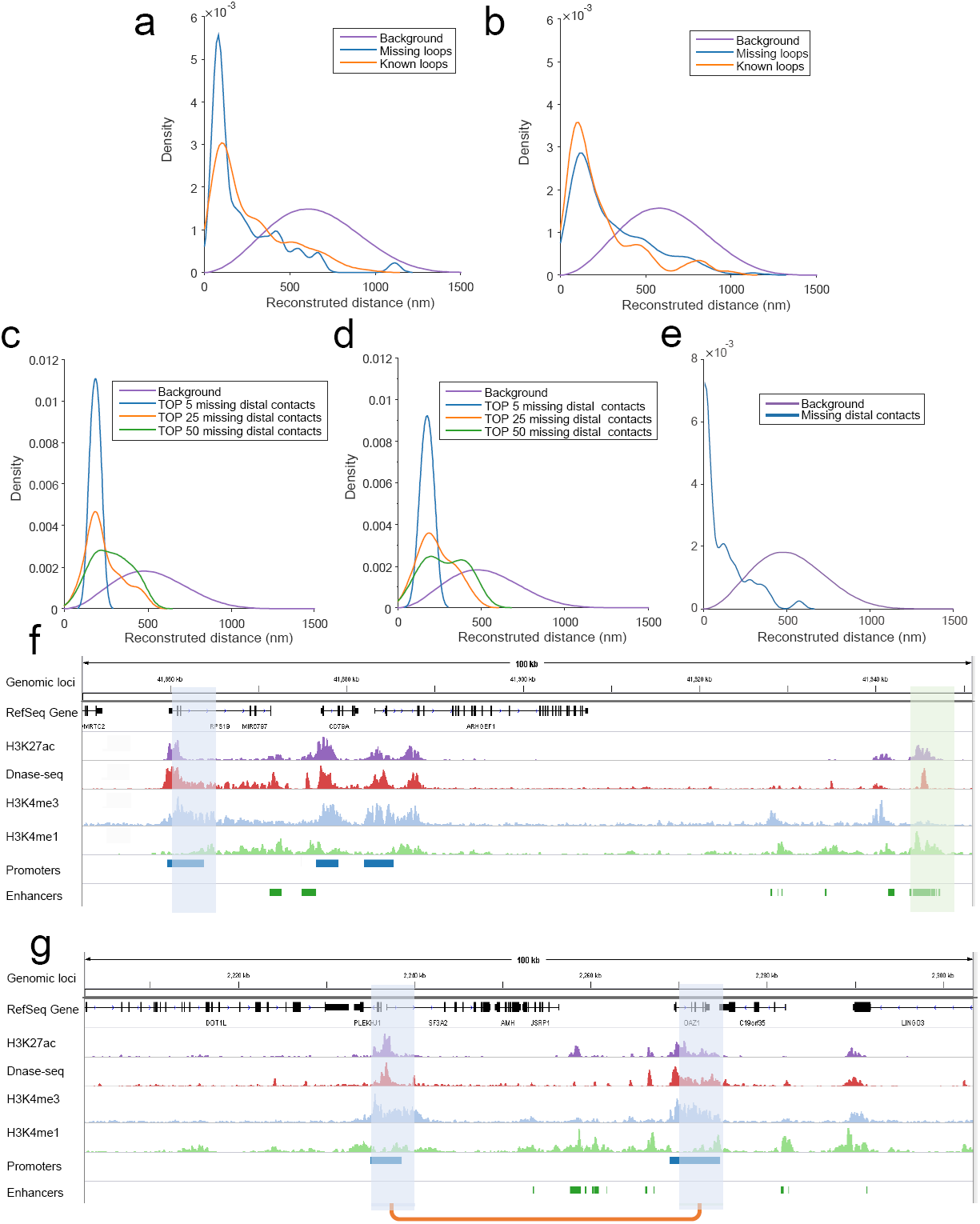
Application of the chromatin structures reconstructed by GEM into the recovery of missing long-range loops or contacts. (**a**,**b**) The recovery results on the missing loops on human chromosome 19 in the GM12878 cell line at 5 kb resolution from the Hi-C data of replicate 1 and replicate 2 [46], respectively. The orange curves represent the distributions of known loops (which were present in the Hi-C data of current replicate), while the blue curves represent the distributions of missing loops (which were missing in current replicate but present in the other replicate). The purple curves show the background distributions, i.e., the distributions of spatial distances in the reconstructed structures. The HiCCUPS algorithm [46] implemented in the Juicer tools [62], with 0.1% FDR, was used to call chromatin loops from Hi-C maps. (**c**,**d**,**e**) The recovery results on the missing promoter-promoter and promoter-enhancer contacts on human chromosome 19, using the chromatin structures reconstructed by GEM based on the promoter-other contacts derived from the capture Hi-C data [47]. The purple curves show the background distributions, i.e., the distributions of all the reconstructed spatial distances (as in Panels **a** and **b**), while the other curves represent the distributions of the promoter-promoter or promoter-enhancer contacts that were missing in the input promoter-other capture Hi-C data [47] but present in an independent Hi-C map (**c**), the promoter-promoter contacts derived from another capture Hi-C data (**d**), or the promoter-enhancer contacts identified by PSYCHIC [48] from an independent Hi-C map [46], all of which were also called the validation Hi-C data. In Panels **c** and **d**, the blue, orange and green curves represent the distributions of the top 5, 25 and 50 missing promoter-promoter contacts which had the highest interaction frequencies in the validation Hi-C data. In Panels **f**, the blue curve represents the distribution of the missing promoter-enhancer contacts in the validation Hi-C data. (**f**,**g**) Two examples on the recovered promoter-enhancer (**f**) or promoter-promoter (**g**) contacts on human chromosome 19 of the GM12878 cell line that were recovered from the chromatin structures reconstructed by GEM from one Hi-C dataset and can be validated by another independent Hi-C dataset. The recovered loops are shown by orange linkers on the bottom, while the connected promoter and enhancers regions (which were annotated using the combination of ENCODE Segway [63] and ChromHMM [64] as in [65]) are shown in blue and green, respectively. Among the lists of chromatin features, H3K27 and DNase-seq signals indicated the active and accessibility states of both ends of chromatin loops, while the states of promoters and enhancers are marked by H3K4me3 and H3K4me1, respectively. All ChIP-seq and DNase-seq data were obtained from the ENCODE portal [66]. The human reference genome GRCh38/hg38 was used.

We also applied the chromatin structures reconstructed by GEM to detect the missing promoter-promoter or promoter-enhancer contacts based on the promoter-other contact map derived from the capture Hi-C technique, a recently developed experimental method to identify promoter-containing chromosome interactions at the restriction fragment level [47]. In capture Hi-C experiments, typically two types of long-range genomic interactions can be observed, i.e., promoter-promoter contacts and promoter-other contacts, depending on whether both ends of the DNA fragments are captured by the promoter regions in the genome. In general, the promoter-other contacts dominate the total number of the interaction frequencies detected by capture Hi-C. Here, we fed all the promoter-other contacts derived from the capture Hi-C data [47] into GEM, and then used the independent Hi-C datasets including conventional Hi-C data [46] and promoter-promoter contacts which were also derived from capture Hi-C experiments [47], to validate those missing long-range genomic contacts recovered by the reconstructed structures. Considering that the distal genomic contacts with more interaction frequencies in Hi-C maps tend to reflect the topological properties of genomic structures with more confidence, here we mainly examined the top 5, 25 and 50 missing promoter-promoter contacts with the highest interaction frequencies in the validation Hi-C data (Fig. 7c,d). In addition, we used the promoter-enhancer contacts identified by PSYCHIC [48] from the conventional Hi-C data [46] to verify the missing distal contacts indicated from the reconstructed structures(Fig. 3e). Our analysis results showed that these recovered promoter-promoter interactions displayed significantly shorter spatial distances than the background of all reconstructed spatial distances (Fig. 7c-e; rank sum test, *P* < 5 × 10^-4^). These results indicated that GEM can be potentially applied to recover the missing long-range genomic interactions caused by the sparsity of the capture Hi-C data.

Careful examination of these missing loops indicated that they were of comparable biological importance to those known chromatin loops, and can also be well supported by the known evidence derived from available chromatin features. For example, the two missing chromatin loops involving promoter-enhancer and promoter-promoter interactions were also consistent with different epigenetic profiles, including chromatin accessibility and histone modification markers H3k27ac, H3k4me3 and H3k4me1 (Fig. 7f,g). In addition, we observed a similar level of the enrichment of functional elements (e.g., H3K27ac, H3k4me3 and H3k4me1 signals, DNA accessible regions, annotated promoter and enhancer regions) in both missing and known chromatin loops (Supplementary Table 1). All these results also demonstrated that the chromatin conformations reconstructed by GEM can provide useful structural templates to recover those missing long-range genomic interactions from the original Hi-C data.

## Discussion and Conclusions

In this work, we have developed a novel manifold learning based framework, called GEM, to reconstruct the 3D spatial organizations of chromosomes from Hi-C interaction frequency data. Under our framework, the 3D chromatin structures can be obtained by directly embedding the neighboring affinities from Hi-C space into 3D Euclidean space, and integrating both Hi-C data and conformational energy. Extensive validations on both simulated and experimental Hi-C data of yeast and human demonstrated that GEM can provide an accurate and robust modeling tool to derive a physically and physiologically reasonable 3D representations of chromosomes.

To our best knowledge, our work is the first attempt to exploit the chromatin structure modeling methods to recover long-range genomic interactions that are missing from original Hi-C data. Here, The ability to recover the missing long-range genomic interactions not only demonstrated a novel extended application of GEM but also provided a strong evidence corroborating the superiority of GEM in terms of physical and physiological reasonability.

Similar to many other computational methods for modeling 3D chromatin structures from interaction frequency data, GEM also faces several technical challenges, e.g., parameter selection and computational efficiency. In GEM, only one parameter (i.e., the coefficient of the energy term *λ_E_*) need to be chosen for an input Hi-C dataset. It can be determined by an automatic parameter tuning method employed in our framework (see “Methods”). In practise, due to the robustness of GEM, the default setting for this parameter often works well for most occasions, which can save the running time required in the parameter selection. Considering that GEM takes a multi-conformation optimization strategy which is usually a time-consuming process, we suggest using a small number of conformations in the ensemble for those tasks with relatively large datasets (e.g., high-resolution Hi-C data) or applications that pay less attention to structural diversity of chromatin structures (e.g., recovery of missing long-range genomic interactions). In principle, more parallel computational schemes can also be employed to further accelerate the optimization process.

## Methods

### Manifold learning framework

Recently, manifold learning, such as t-SNE [32], has been successfully applied as a general framework for nonlinear dimensionality reduction in machine learning and pattern recognition [30, 33–35]. It aims to reconstruct the underlying low-dimensional manifolds from the abstract representations in the high-dimensional space, that is, it uncovers the intrinsic low-dimensional manifolds which preserve the local neighbourhoods of high-dimensional data (Supplementary Fig. 1).

In practice, many high-dimensional data of interest lie on the structures that are intrinsically embedded from a low-dimensional manifold. For example, in computer vision, images of faces can be regarded as points in a high-dimensional vector space, in which each dimension corresponds to the brightness of every pixel in the image. All the images can be considered to lie on an intrinsic 3D manifold parameterized by two pose variables and and an azimuthal lighting orientation angle [34]. In our chromatin structure modeling problem, the meaningful spatial organizations of chromosomes can be interpreted as the geometry of manifolds in 3D Euclidean space. The Hi-C interaction frequency data can be regarded as a specific representation of the neighboring affinities reflecting the spatial arrangements of genomic loci, which is intrinsically determined by the underlying manifolds embedded in Hi-C space. Thus, manifold learning can be used here to uncover the meaningful geometry of manifolds in low-dimensional space based on a process of neighborhood embedding, which preserves the local neighborhood of genomic loci in Hi-C space.

### Modeling the conformational energy of chromatin structures

According to our known biophysical knowledge of a polymer model [49–51], the physical potential of a chromatin conformation can be described by an energy function *E*^(*m*)^ consisting of three terms, including the stretching energy *E*_*stretch*_, the bending energy *E*_*bend*_ and the excluding energy *E*_*exclude*_, that is,

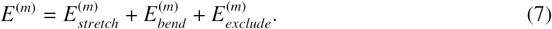

First, the stretching energy term *E*_*stretch*_ accounts for the stretching resistance of chromatin fibers, which is defined by the following equation

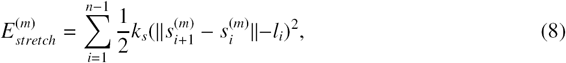

where *k*_*s*_ stands for the spring constant characterizing the chromatin stiffness, and *l*_*i*_ stands for the equilibrium length of the *i*-th segment in the modeled chromatin structure. Since the sequence length for each pair of adjacent genomic loci is known, their corresponding distance *l*_*i*_ can be generally derived based on the packing density, which is usually assigned with 130bp/nm [36, 52] in approximate 10 kb resolution and can be computed based on a 1/3 power-law relationship between spatial distances and corresponding genomic distances in other resolutions (which can be derived mainly based on the 3D FISH data [13] or the fractal globule model [53]).

Second, the bending energy term *E*_*bend*_ accounts for the bending potential of chromatin fibers [54], which is defined by the following equation,

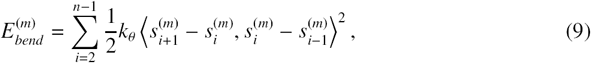

where *k*_θ_ denotes the bending constant and ⟨·⟩ denotes the angle between two adjacent segments.

Third, the excluding energy term *E*_*stretch*_ accounts for the inter-particle repulsive potential. It takes the form of the repulsive part of the Lennard-Jones potential [55, 56], that is,

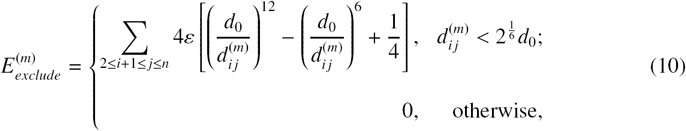

where 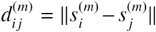 stands for the Euclidean distance between 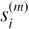 and 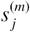, ε is a Lennard-Jones energy parameter, and *d*_0_ is the distance threshold within which the repulsive force is zero.

### Optimization of the objective function

We use gradient descent to minimize *C* in Equation (4). In particular, the gradient of *C* is calculated as follows

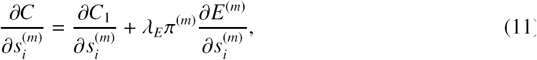

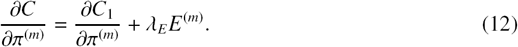

The gradient of *C*_1_ with respect to the coordinates *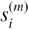* in 3D Euclidean space is given by

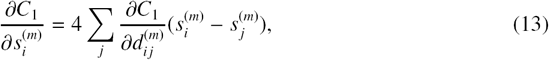

where 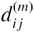 stands for the Euclidean distance between 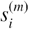 and 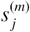, and

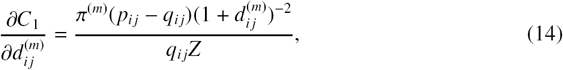

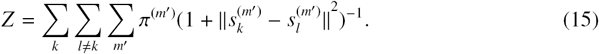

From the term 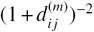 in the above gradient, we can also see that two neighboring nodes are not likely to be modeled by widely separated points. Also, π^(*m*)^ is defined as follows to satisfy the definition of probability, that is,

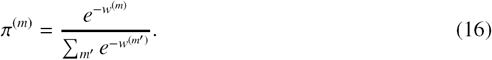

Then we can obtain a new version of the gradient

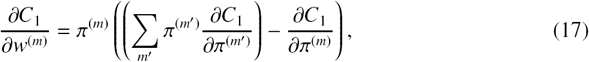

where

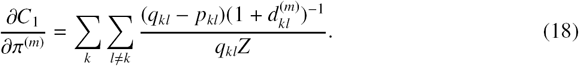

The gradient of *E*^(*m*)^ with respect to *s*^(*m*)^ includes three parts, that is,

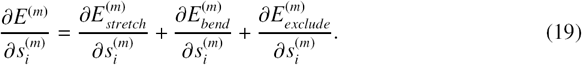

The first part is given by

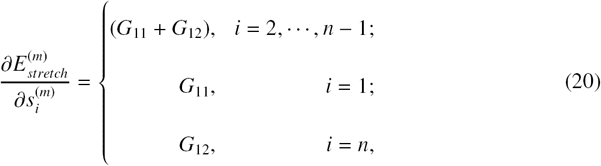

where

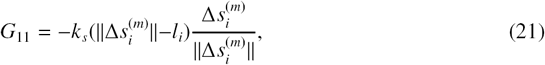

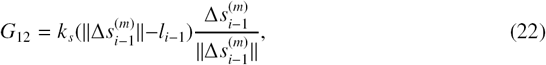

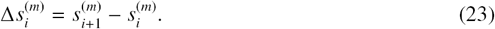

The second part is given by

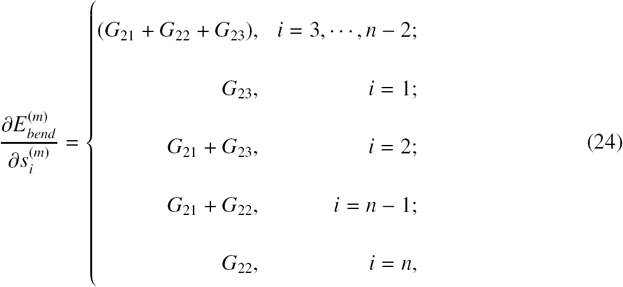

where

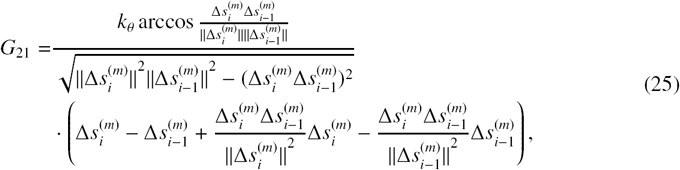

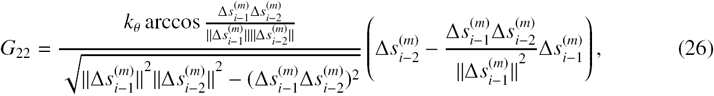

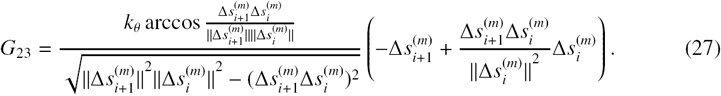

The third part is given by

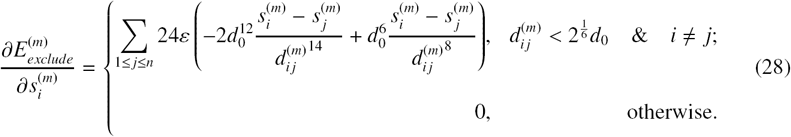

Based on the above derivations, we can develop an adaptive gradient descent method to solve the optimization problem in Equation (4). In particular, the learning rates for π^(*m*)^ and 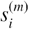 can be changed constantly during different stages to accelerate the optimization process. However, there may exist a “sinking” problem in such an optimization strategy, if the random initialization process leads to a huge difference of goodness between a pair of conformations. In such a case, if the learning rate for π^(*m*)^ is too large, π^(*m*)^ of those structures that deviated largely from random initialization will sink at zero rapidly, and these conformations will not be considered during the downstream optimization process, mainly due to the vanishing gradient, although they can still converge to proper solutions if a sufficient number of optimization iterations are performed.

To address this problem, we could employ two strategies during the optimization process, i.e., two-stage optimization and asynchronous starting. The first strategy is to divide the whole optimization procedure into two stages, including average-structure optimization and multiconformation optimization. In particular, we first compute an average structure using a single-conformation version of GEM. Then, this average structure is used as an initial structure for the second-stage optimization, which is accomplished through a multi-conformation version of GEM. Initialization from such a pre-computed structure that is not so far away the final solution provides a beneficial guidance for optimization. More importantly, the goodness scores of the initial conformations in the second-stage optimization do not has extremely large variance, which thus can prevent the aforementioned sinking problem. The second strategy that we could use is to delay the update of the learning rates of π^(*m*)^ in the second-stage optimization if the goodness scores of the initial conformations remain considerably different. Under such a strategy, it is unlikely that the sinking problem will occur, after a number of iteration steps to optimize the conformations with relatively fixed weights. In practice, we found that the first strategy is often sufficient enough to prevent the sinking problem.

### The convergence and parameter selection of GEM

We examined the convergence of the optimization procedure employed in GEM, which is a two-stage optimization scheme including average-structure optimization and multi-conformation optimization. As shown in Supplementary Fig. 5a, both optimization stages converged successfully. In the first optimization stage, which aimed at computing the average structure, the cost function descended rapidly at the beginning. After 2000 iterations, the cost function began to converge, indicating a stable average structure was reached. After that, the second optimization stage was performed to obtain multiple conformations. Probably because the average structure had been determined, the cost function only descended slightly in this stage. Overall, the second optimization stage converged after approximate 20000 iterations.

In our model, the only parameter, the coefficient of the energy term *λ_E_*, determines a trade-off between the fitness of the spatial constraint derived from Hi-C data and structural feasibility measured in terms of conformational energy. The parameter *λ_E_* can be decided by the users according to their emphasized aspects. Alternatively, this parameter can be determined by the following two automatic methods.

First, inspired by previous studies [19, 21, 57], we can use a Bayesian approach to determine the proper value of the coefficient of the energy term, *λ_E_*, based on extra priori knowledge, such as the volume of a chromosome obtained by direct experimental observations or estimated by indirect experimental observations (e.g., DNA density [58]). The goal is to select a proper value of *λ_E_* that best interprets both the input Hi-C data and the observation about the volume of a chromosome. The posterior probability Pr(*λ_E_*|*H*, *V*) of the coefficient *λ_E_* given Hi-C data *H* and the volume of a chromosome *V* can be derived according to Bayes’ theorem, i.e.,

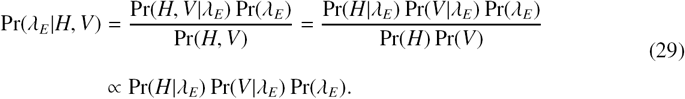

Based on the maximum a posteriori estimation of *λ_E_*, we define a Bayesian score to evaluate the parameter *λ_E_* for our model. We assume that the prior distribution of *λ_E_* is uniform, and thus Pr(*λ_E_*) can be considered constant and is not necessary to be included in the Bayesian score. The final optimized value of the KL divergence *C*_1_ mentioned in Equation (4), which is dependent on *λ_E_* during the optimization process, measures the degree of mismatch (ranging from 0 to 1) between structures calculated with *λ_E_* and the Hi-C data. Here, we use 1 - *C*_1_ to define Pr(*H*|*λ_E_*). In addition, the mismatch between the volume of a computed structure *v*’ (which is also dependent on *λ_E_*) and its real volume *v* can be measured by the relative error ratio. Here, we use the inverse of this relative error ratio to define Pr(*V*|*λ_E_*). To sum up, the Bayesian score is defined as,

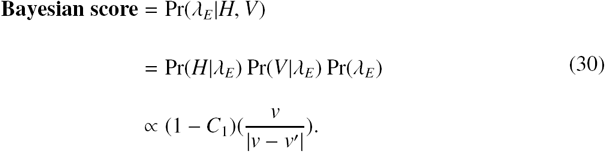

Second, if we do not have any priori knowledge, we can transform the parameter selection into a multi-criteria decision problem and use TOPSIS [59] to obtain the best estimate of parameter *λ_E_*. In this setting, each value of different *λ_E_* is regarded as a decision that is evaluated by two criteria *C*_1_ and *C*_2_.

In our study, we employed the Bayesian approach to perform parameter selection. We computed the Bayesian scores with respect to a wide range of the coefficient of energy term *λ_E_* (Supplementary Fig. 5b) and then chose *λ_E_* = 5 × 10^-12^ which had the maximum Bayesian score, as the most reasonable parameter for human chromosome 14 (which is marked by the orange dashed line in Supplementary Fig. 5b). In practice, we only need to select a rough range for *λ_E_* because in general the changes of *λ_E_* within the same order of magnitude have little influence on the performance of our model, measured by the Pearson correlation between experimental and reconstructed Hi-C data in the 10-fold cross-validation procedure (Supplementary Fig. 5c).

### Generation of simulated Hi-C data

The simulated Hi-C data were generated according to the following procedure. At the beginning, we applied the Brownian simulation method to generate a set of random chromatin conformations, each of which imitated a real chromatin conformation in Hi-C experiments. Let *N*_*c*_ denote the total number of cells. Then we mimicked the experimental Hi-C protocol to obtain the simulated Hi-C interaction frequency map. First, the restricted sites along the synthetic chromatin conformations were chosen randomly and then cleaved by the restriction enzymes. Second, as in the study by Trussart et al. [20], we used a Gaussian process to generate the genomic interactions between restriction sites. In particular, let *P*_*m*_ and σ denote the scaling factor (i.e., the maximum interaction probability) and the standard deviation of the Gaussian function describing the relationship between the probability of generating the interaction and the spatial distance between a pair of genomic loci. Due to experimental uncertainty, not all the interactions between restriction sites can be captured by Hi-C experiments. Here, we used the trapping rate *α_t_* to model such experimental uncertainty, this is, with probability *α_t_*, an occurred interaction is observed between restriction sites, otherwise it is missed with probability 1 - *α_t_*. Overall, the simulation process for generating a synthetic interaction frequency dataset can be determined by parameters (*α_t_*, *P*_*m*_, σ, *N*_*c*_).

### The 10-fold cross-validation procedure

The Hi-C data of a chromosome were randomly divided into 10 roughly equal-sized subsets. Nine of them were selected as training data and input into GEM to compute the chromatin structures. Based on the latent function between interaction frequencies and spatial distances between genomic loci derived by GEM (see the dashed box in Fig. 1), the modeled chromatin structures can also be used to obtain the reconstructed or predicted Hi-C map. Then the remaining subset was held as test data to assess the accuracy of the modeled conformations by comparing the original Hi-C map to the reconstructed Hi-C map. Such a process was performed 10 folds, and the average result was used to evaluate the final modeling performance.

## Data availability

The GEM model and the analysis data files can be downloaded from https://github.com/mlcb-thu/GEM. The Hi-C data of yeast can be downloaded from Duan et al. [14] (http://noble.gs.washington.edu/proj/yeast-architecture/sup.html). The Hi-C data of human used for model validation can be downloaded from NCBI GEO GSE18199 [7] (https://www.ncbi.nlm.nih.gov/geo/query/acc.cgi?acc=GSE18199) and NCBI GEO GSE48262 (https://www.ncbi.nlm.nih.gov/geo/query/acc.cgi?acc=GSE48262), and the normalized version can be downloaded from Yaffe et al. [60] (http://compgenomics.weizmann.ac.il/tanay/?page_id=283). The Hi-C data and capture Hi-C data of human used for the recovery test are available in NCBI GEO GSE63525 (https://www.ncbi.nlm.nih.gov/geo/query/acc.cgi?acc=GSE63525) and ArrayExpress E-MTAB-2323 (http://www.ebi.ac.uk/arrayexpress/files/E-MTAB-2323/), respectively.

## Acknowledgments

This work was supported in part by the National Natural Science Foundation of China Grant 61472205 and China’s Youth 1000-Talent Program, the Beijing Advanced Innovation Center for Structural Biology and NCSA Faculty fellowship 2017. T.K. is a member of the Israeli Center of Excellence (I-CORE) for Gene Regulation in Complex Human Disease (no. 41/11) and the Israeli Center of Excellence (I-CORE) for Chromatin and RNA in Gene Regulation (no. 1796/12), and is also supported by the Israel Science Foundation (grant no. 913/15). We thank the members from Prof. Cheng Li’s group and Prof. Michael Zhang’s group for helpful discussions.

## Author contributions

G.Z. and J.Z. conceived the research project. J.Z. supervised the research project. G.Z., W.D., J.P. and J.Z. proposed the GEM model. G.Z. preprocessed raw data, designed the details of GEM model, developed optimization process and performed model evaluation. G.Z., H.H., T.K. and J.Z. accomplished the task of recovering the missing loops and contacts. G.Z., R.M., S.Z., J.P. and J.Z. fine-tuned the mathematical part of the methods. G.Z., W.D., H.H., R.M., S.Z., J.Y., T.K. and J.Z. discussed the results and performed the computational analyses. G.Z., H.H. and J.Z. wrote the manuscript with the help from other authors.

## Supporting Material

**Supplementary Figure 1:**
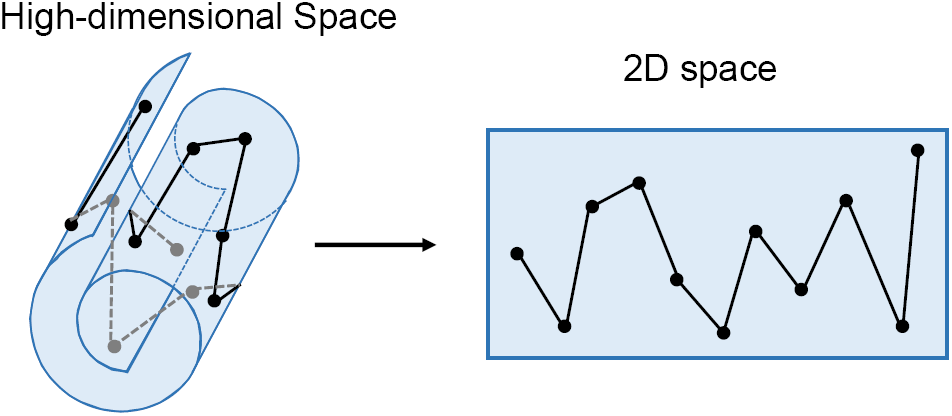
A schematic illustration of manifold learning. In high-dimensional (3D for example) space, the data points lie on a “Swiss roll” structure. After embedding the data points from high-dimensional space into low-dimensional (2D for example) space by manifold learning, the intrinsic manifold is uncovered.

**Supplementary Figure 2:**
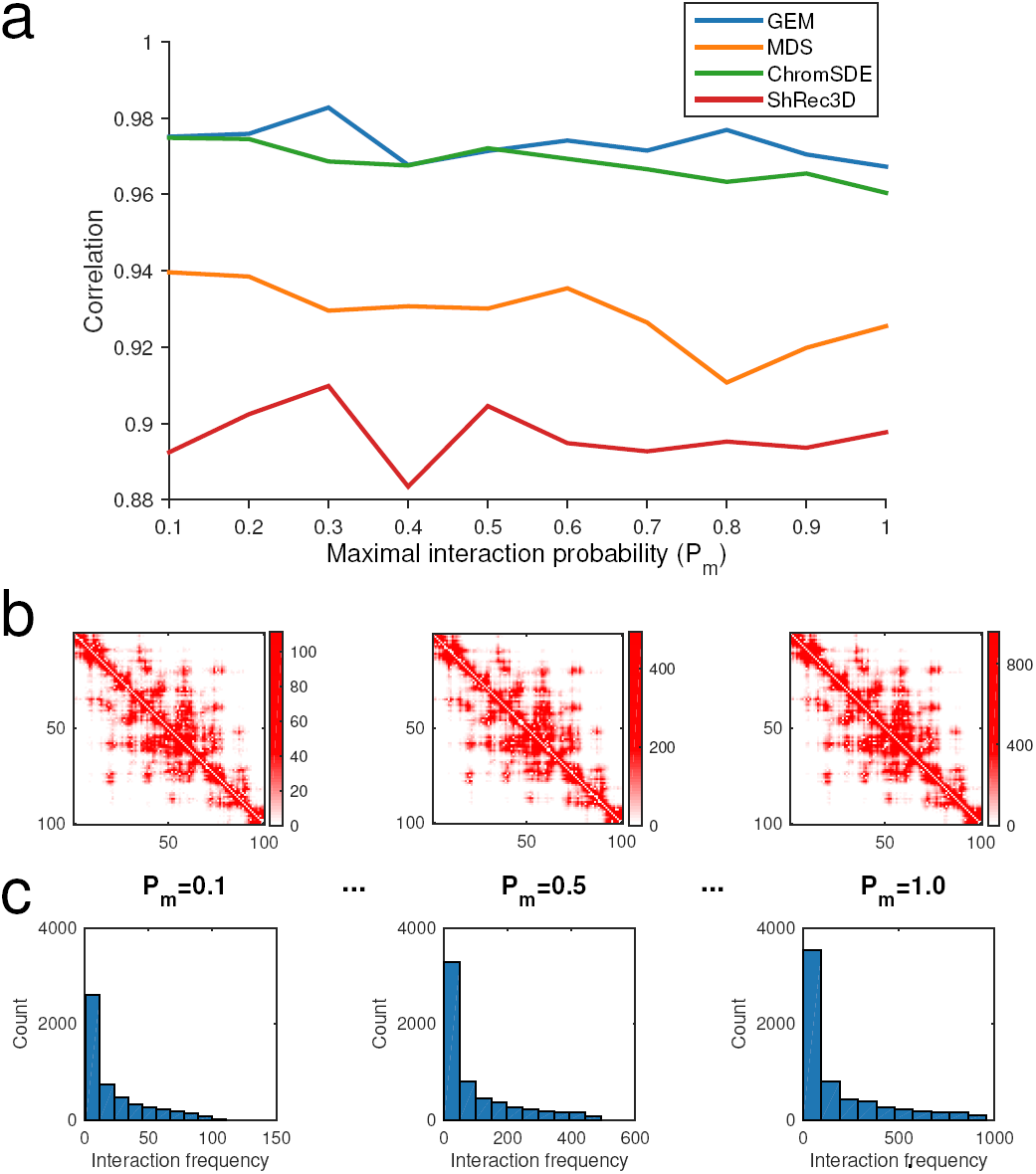
The validation results on the simulated Hi-C data, which were generated according to different settings of the maximum interaction probability *P*_*m*_ (see “Methods”). (**a**) The comparisons of Pearson correlations between GEM and other modeling methods, including the MDS based model [29], ChromSDE [17] and ShRec3D [18]. (**b**) and (**c**) show the typical examples of the simulated Hi-C maps and the corresponding distributions of the reconstructed interaction frequencies as *P*_*m*_ increases, respectively.

**Supplementary Figure 3:**
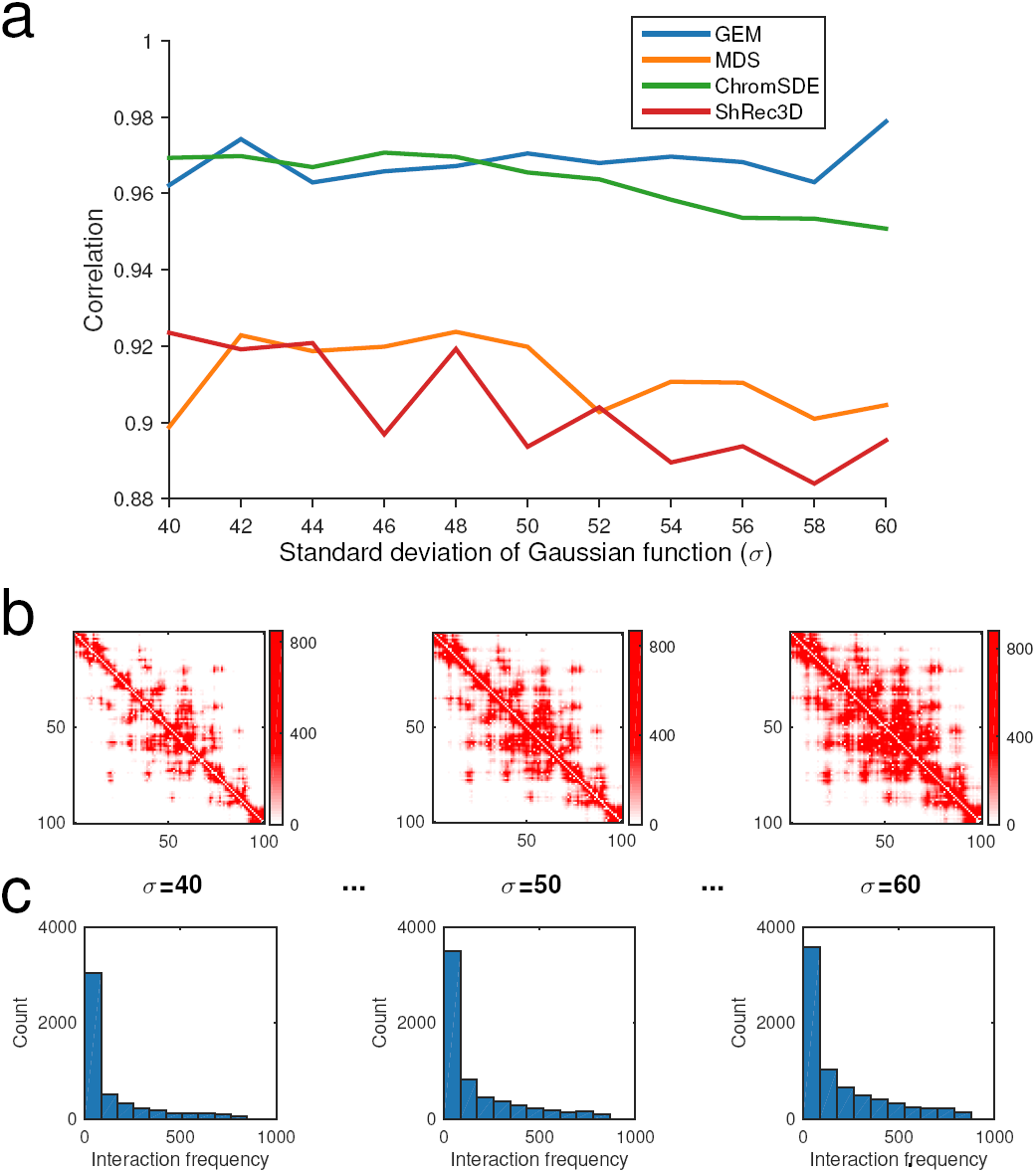
The validation results on the simulated Hi-C data, which were generated according to different settings of the standard deviation of Gaussian function σ (see “Methods”). (**a**) The comparisons of Pearson correlations between GEM and other modeling methods, including the MDS based model [29], ChromSDE [17] and ShRec3D [18]. (**b**) and (**c**) show the typical examples of the simulated Hi-C maps and the corresponding distributions of the reconstructed interaction frequencies as σ increases, respectively.

**Supplementary Figure 4:**
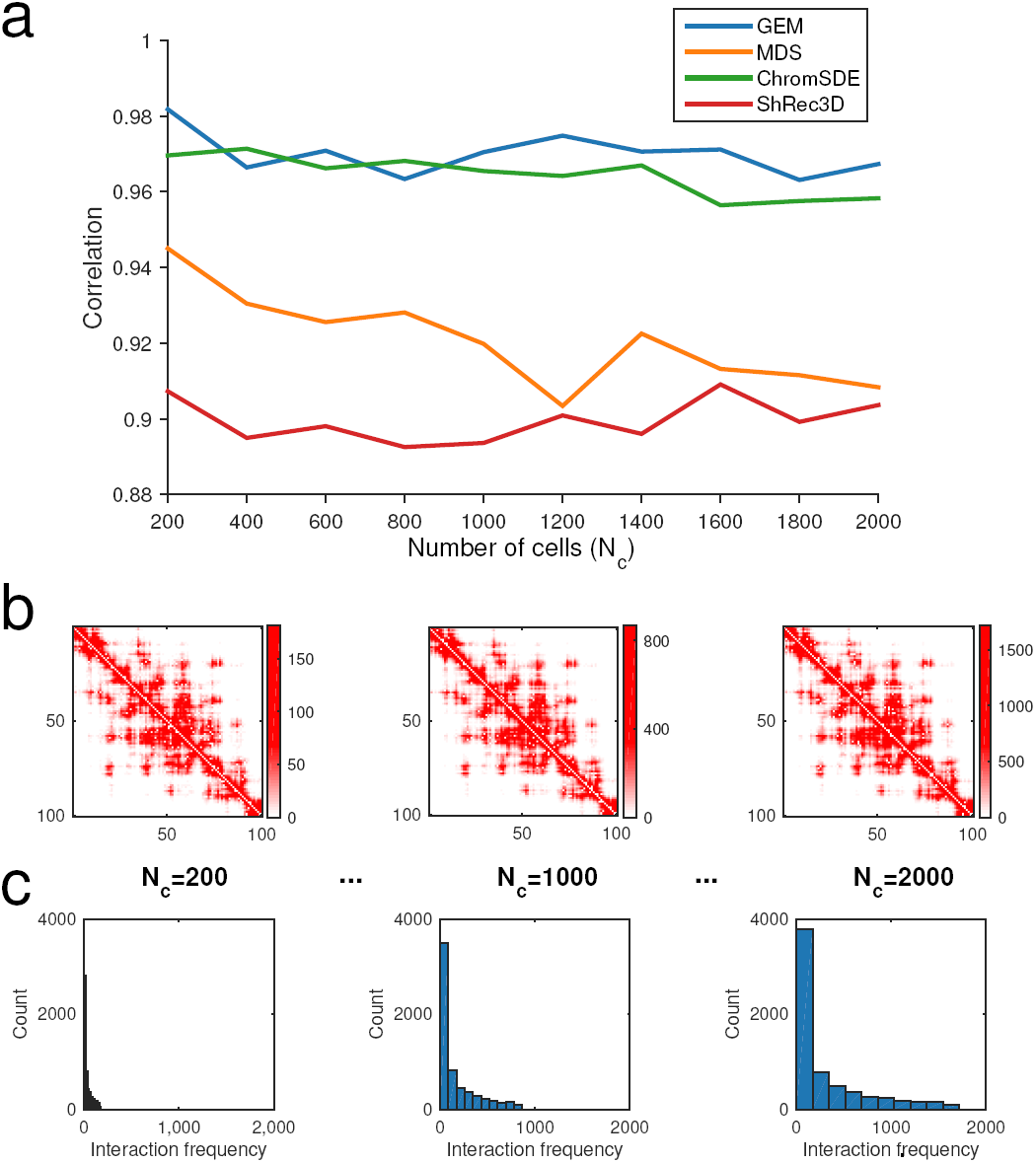
The validation results on the simulated Hi-C data, which were generated according to different settings of the number of cells *N*_*c*_ (see “Methods”). (**a**) The comparisons of Pearson correlations between GEM and other modeling methods, including the MDS based model [29], ChromSDE [17] and ShRec3D [18]. (**b**) and (**c**) show the typical examples of the simulated Hi-C maps and the corresponding distributions of the reconstructed interaction frequencies as *N*_*c*_ increases, respectively.

**Supplementary Figure 5:**
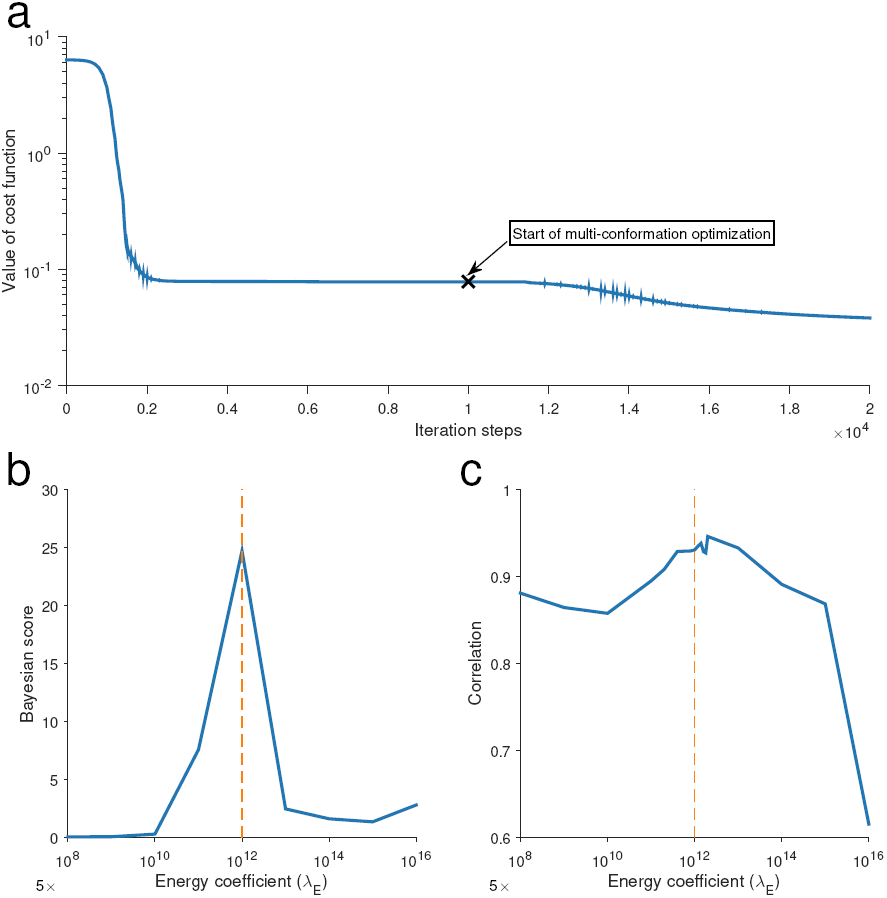
The convergence and parameter selection of GEM. Here, human chromosome 14 at a resolution of 1 Mb was used as an example. (**a**) The convergence results on the cost function *C* in Equation (4). The first-stage optimization (i.e., average-structure optimization) took about 1000 iterations and stopped at the position marked with black cross, which denotes the start of the second-stage optimization (i.e., multi-conformation optimization), which also took about 1000 iterations. (**b**) The Bayesian score as a function of the coefficient parameter *λ_E_* that weighs the conformational energy term. The value of *λ_E_* (5 × 10^-12^) with the maximum Bayesian score was used in GEM, which is marked by the orange dashed line. (**c**) The 10-fold cross-validation results (evaluated in terms of the Pearson correlation between experimental and reconstructed Hi-C data) as a function of the coefficient parameter *λ_E_* for the conformational energy term. The orange dashed line indicates the value of *λ_E_* used in GEM.

**Supplementary Table 1:**
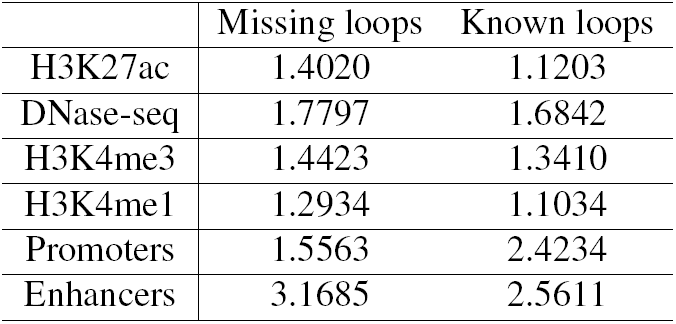
The enrichments of the functional elements (compared with the background) in both missing and known chromatin loops on human chromosome 19.

|  | Missing loops | Known loops |
| --- | --- | --- |
| H3K27ac | 1.4020 | 1.1203 |
| DNase-seq | 1.7797 | 1.6842 |
| H3K4me3 | 1.4423 | 1.3410 |
| H3K4me1 | 1.2934 | 1.1034 |
| Promoters | 1.5563 | 2.4234 |
| Enhancers | 3.1685 | 2.5611 |

